# Range shifts of Eastern South American mangroves in a changing climate

**DOI:** 10.64898/2026.06.11.731615

**Authors:** Miguel P. Pereira-Romeiro, Gustavo M. Mori, Flávia M. D. Marquitti

## Abstract

As climate changes, habitat suitability for multiple taxa are also expected to change. In recent years, mangrove latitudinal range expansion has been linked to increasing temperatures and reduced freezing in temperate regions, happening mainly through encroachment into saltmarshes. The range limit of mangrove forests in Eastern South America has not seen drastic changes in the last four decades, despite trends of increased temperature and the seemingly favorable direction of the Brazilian Current. Here, we investigate if and how the distribution of South-Atlantic mangrove forests may respond to different scenarios of climate change. To do this, we combine ecological niche modelling with propagule dispersal simulations to understand the roles of climate and ocean currents in defining the austral limits of South-Atlantic American mangroves. Our results indicate that minimum sea surface temperature strongly constrains habitat suitability beyond the current distribution of mangroves (28°28’68” S), while dispersal processes heavily limit propagule stranding beyond 35° S. The Brazil–Malvinas currents confluence zone creates steep temperature gradients and an oceanographic barrier that makes the latitudinal expansion of mangroves unlikely in this region, even in future scenarios of heating climate. We found no evidence of current nor future poleward expansion of mangroves, but total mangrove area has increased in Brazil over the last decades, likely due to landward migration, but anthropogenic interference and urban expansion may restrict this process, leading to coastal squeeze. Under scenarios where both landward and poleward migration are limited, South American mangroves may face increasing vulnerability, with potential impacts on the several ecological, biogeochemical and social cycles they support. Our results contribute to leading hypotheses of climate restriction and shed light on the role of ocean currents in South America, helping to explain why the poleward expansion reported in other regions has not yet been observed in the South-Atlantic mangrove range limit.

## Introduction

As climate changes, habitat suitability for multiple taxa are also expected to change. With shifts in climatic regimes, temperate regions may experience a reduction in extreme cold events and increased temperatures. This change in climate may allow the range expansion of tropical or cold-limited taxa into new areas, which may come at the expense of the distribution of temperate adapted organisms (Osland et al., 2021). Because of the implications of such widespread shifts, understanding the impacts of climate change in the distribution of species and ecological systems is rapidly becoming one of the main pressing issues in modern ecology (Osland et al., 2021).

Coastal wetlands and their resident species are considered to be especially sensitive to climate driven changes, as they occupy the land-sea interface and may suffer from threats coming from these different environments (Alongi, 2015). Mangrove forests offer a form of coast protection, as they help mitigate erosion and tropical storms, and their preservation is deemed as a valuable nature-based solution for coastal resilience (Barbier et al., 2011; Lee et al., 2014; Walters et al., 2008). Despite this, these ecosystems are also at risk, as they are not only vulnerable to sea-level rise (Osland et al., 2016; Saintilan et al., 2020) and anthropogenic impacts (Goldberg et al., 2020; Hagger et al., 2022; Heck et al., 2024; Vanin et al., 2025a), but also to the effects of macroclimatic drivers, such as changes in temperature, precipitation and storm regimes (Friess et al., 2022; Osland et al., 2016). For instance, mangroves are generally limited by low air/sea temperatures and rainfall, but also extreme events such as freezings and hurricanes, which may lead to forest dieback (Adame et al., 2021; Devaney et al., 2021; Hickey et al., 2017; Osland et al., 2017; Saintilan et al., 2014). These climatic restrictions make the distribution of mangrove forests relatively well defined latitudinally, occurring mainly in tropical and subtropical climates. Saltmarshes, by contrast, seem to be less constrained by climate than mangroves, occurring predominantly in temperate regions and high latitudes (Friess et al., 2012).

Paleobiogeographical evidence suggests that the distribution of mangrove forests have changed in response to shifts in the macroclimatic regime of the planet (Osland et al., 2017; Srivastava & Prasad, 2019), reaching as far north as 72° in the Eocene (Suan et al., 2017). In recent years, mangrove range expansion has occurred mainly through encroachment into saltmarshes, which has been linked mainly to increasing temperatures and reduced freezing in temperate regions (Friess et al., 2022; Hickey et al., 2017; Saintilan et al., 2014; A. C. Ximenes et al., 2021). Despite this general trend of expansion, pinpointing how the distribution of mangroves might respond to climate change is not trivial, as the temperature and precipitation thresholds associated with latitudinal range limits of mangrove forests vary across the world (Quisthoudt et al., 2012; Simard et al., 2019). Additional variability in responses may arise when considering the role of coastal processes and ocean currents, as most mangroves genera depend on them for propagule dispersal (Van Der Stocken et al., 2019). Thus, beyond temperature and precipitation related changes, processes that control the dispersal potential of mangroves will likely influence responses in mangrove distribution, especially near range limits, as they may favor or restrict dispersal of propagules to new areas. For example, in North America, poleward expansion of mangroves in the west coast is limited mainly by dispersal and not by temperature (Cavanaugh et al., 2024), whereas populations on the east coast seem to be more climate limited (Enes Gramoso et al., 2026).

In South America, the latitudinal limits of mangrove occurrence vary vastly between the Pacific and Atlantic coasts. The Pacific limit occurs at 5°30’ S and is mainly attributed to the aridity of the coast and intense upwelling (Clüsener & Breckle, 1987; Saintilan et al., 2014; A. C. Ximenes et al., 2025). In the Atlantic side of the continent, the southernmost occurrence is at Santo Antônio Lagoon (28°28’ S), where mangrove trees present reduced height and structural development (Soares et al., 2012). This stunted growth is possibly an indicative of climatic limitation for mangrove development, related to low temperatures but not aridity, as precipitation does not seem to be an issue for mangrove development in the Brazilian coast (Schaeffer-Novelli et al., 1990; Soares et al., 2012; A. Ximenes et al., 2016). The Atlantic side of South America is also characterized by the Brazilian current that originates near the equator and runs southwards along the coast of the continent. Despite the seemingly favorable direction of the Brazilian Current, the range limit of mangrove forests in this coast has not seen drastic changes in the last four decades (Schaeffer-Novelli et al., 1990; Soares et al., 2012), with a ∼10ha mangrove area expansion between 2003 and 2019 (Cohen et al., 2020). Because of these characteristics, we investigated the roles of climate and ocean currents as potential limiting factors in defining the austral limits of South-Atlantic American mangrove forests. We also aimed to assess if and how the distribution of South-Atlantic mangrove forests may respond to different scenarios of climate changes. We begin by estimating environmental suitability for mangrove occurrence both in current and future (2050) climates along the eastern South Atlantic coast. Then, using propagule dispersal simulations, we estimate connectivity between contemporary mangrove forests and estuaries beyond their current range, in order to identify potential colonizing events of new areas and understand the overall dispersal dynamics of current mangrove forests. By explicitly incorporating dispersal constraints into environmental suitability models, we aimed to bridge the gap between the fundamental and realized niches of mangroves, enabling more robust predictions of potential distribution shifts under climate change.

## Methods

### 1. Study area and sample points

Schaeffer-Novelli et al. (1990) proposed that the Brazilian coast can be divided into eight distinct segments defined by mangrove climatological, hydrographic and oceanographic drivers. Because our goal was to investigate potential latitudinal range shifts of mangrove forests, we focused on segments V to VIII (05°08’ S - 32°35’ S) (Fig. 1), which encompass the current southern distribution limit of mangroves and are considered to form a single population (Madeira et al., 2023). This choice is further supported by large-scale oceanographic dynamics (Results). The South Equatorial Current splits when it meets the South American continental shelf, feeding both the North Brazil Current and the Brazil Current, which diverge and flow in opposite directions. This pattern limits propagule exchange between northern segments (I - IV) and southern segments (V - VIII). As a result, potential poleward expansion would likely depend on propagules originating from segments V to VIII.

**Figure 1.**
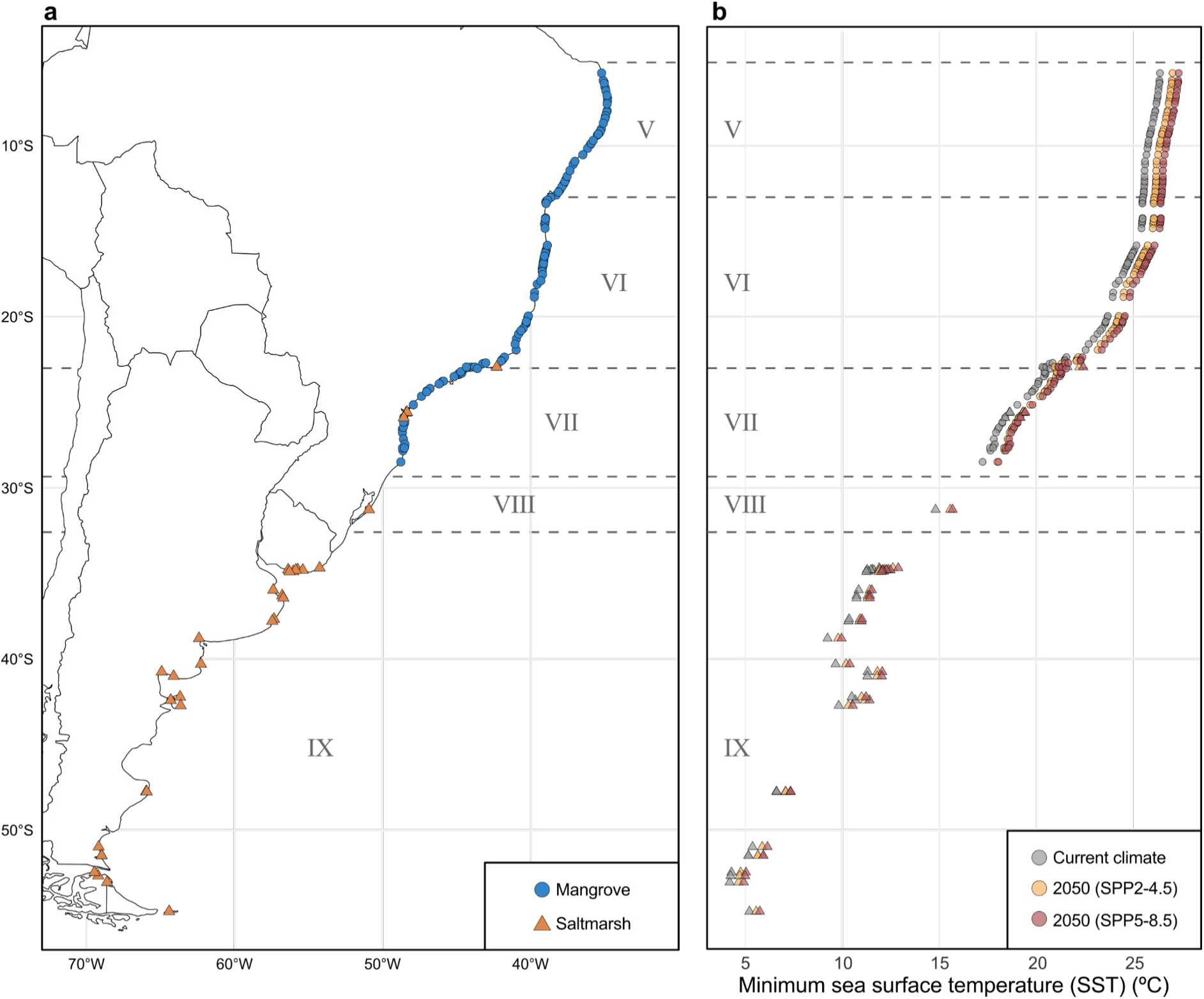
Mangrove and saltmarsh sites (a). Current and future (2050) minimum sea surface temperature (x-axis) for each site (y-axis) (b).

We used the Google Earth Engine code editor to filter mangrove patches. To do this, we isolated pixels classified as mangroves in these segments from the MapBiomas 8 collection (30m resolution) (Souza et al., 2020) and then used a square shape kernel (10 pixels) to identify neighboring mangrove pixels and split the total mangrove area into different patches. For this, each mangrove pixel had its neighboring pixels checked in a square spanning 10 pixels in each direction (approximately 300 m). If any mangrove pixels were within that range, they were considered to be of the same patch. Patches were then labeled and exported as a raster file with 100m/px resolution. In R 4.2.2 (R Core Team, 2024), using the *terra* v.1.7.78 package (Hijmans, 2020), we calculated the area of each patch. Then, we transformed each patch into a polygon and defined a centroid for each one. For each mangrove centroid, we did two transformations: we (I) snapped each centroid to the closest point in the coastline, using a high resolution shapefile of Brazil (South, 2011) and the *snap* function in QGIS v.3.34.10 (Nyall Dawson et al., 2024); then, in R, we (II) projected the snapped points 5km into the ocean by using a bisector line that crossed the coastline at the snapped point (Fig. S1). For saltmarshes, we obtained data from McOwen et al. (2017), which is the product of a thorough review of the distribution of saltmarshes from across the globe. We extracted the geographical location of all saltmarshes in the Atlantic coast of South America and snapped these points to the closest point in the coast, using the same process we did for mangrove centroids, but did not project these points into the ocean. In total, we obtained 122 mangrove and 34 saltmarsh points, and each version of these points (original; snapped to the coast or projected into the ocean) was used for subsequent analyses. Snapped points with identification, geographical location and vegetation can be found in Table S3.

### 2. Environmental niche modelling (ENM)

To predict changes in the distribution of mangrove forests, we first used an ecological niche model (ENM). Low temperatures are considered a general barrier for mangrove growth and may render propagules unviable (Van Der Stocken et al., 2019). Aridity also limits mangrove growth (Adame et al., 2021), however, precipitation in the studied area is generally high (A. Ximenes et al., 2016; A. C. Ximenes et al., 2021) and therefore is arguably not limiting to mangrove development. Thus, we decided to use variables related to minimum sea surface temperature and minimum air temperature as predictor variables in our model (Cavanaugh et al., 2024; Enes Gramoso et al., 2026; Raw et al., 2023). Historical climate data we collected referred to 2010 to 2020. Future data referred to 2040 - 2060 and to two Shared Socioeconomic Pathway (SSP) scenarios of climate change: SSP2-4.5 and SSP5-8.5.

We obtained climate data from WorldClim version 2.1 (Fick & Hijmans, 2017), a database of global weather and climate data in high spatial resolution. Beyond raw climate data, WorldClim also makes available 19 bioclimatic derived from monthly temperature and rainfall variables that represent trends, seasonality and limiting environmental conditions. We downloaded monthly temperature (minima and maxima) and precipitation data from 2010 to 2020 and, using the *dismo* v.1.3.16 package (Hijmans et al., 2010) in R 4.2.2 (R Core Team, 2024), generated the 19 bioclimatic variables for this period. In order to use variables that are biologically sound with mangrove limiting factors, we selected the minimum temperature of the coldest month (BIO6).

WorldClim also provides future climatic data based on multiple different global circulation models (GCMs). In order to determine a suitable GCM for our interest area, we utilized the *chooseGCM* v.1.2 package (Esser et al., 2025). This package is designed to help select GCMs based on the variation of predictor variables in the area of interest. Future data from available GCMs were downloaded from WorldClim 2.1 at 2.5 arcmin resolution (∼21 km2 at the equator) (Fick & Hijmans, 2017). After selection, we used data from the CanESM5-CanOE GCM (Christian et al., 2022).

For sea surface temperature, we obtained data from Bio-Oracle v3.0 (Assis et al., 2024), a global environmental dataset available at a spatial resolution of 0.05° and a temporal resolution of 10 decadal steps. For sea surface temperature (SST), we used the long-term average of the yearly minima (e.g., the average temperature of the coldest month in the sampled period). To avoid variable redundancy, we calculated the correlation coefficient between BIO6 and SST and found that they were highly correlated (*r* = 0.99, *p* < 0.01), thus, we retained only SST as a predictor for the ENM.

To model the potential distribution of mangrove forests, we used the *biomod2* v.4.2.5.2 package (Thuiller et al., 2012) in R 4.2.2 (R Core Team, 2024). This model works by making predictions of species distribution based on presence and (if available) absence data points and climatic variables. We defined the snapped points obtained in the first step (see *1. Study area and sample points*) as presence data. Historically, mangrove forests have been extensively sampled, either through field work or satellite imaging (Jia et al., 2023), which makes their spatial distribution well recorded. Absence or non-detection data is usually difficult to obtain, which may make modelling species distributions not as straightforward (Franklin, 2023). Long stretches of coast without forests are not uncommon and are caused by geomorphological rather than climatic factors, such as sandy beaches and/or dunes, which are not adequate for propagule recruitment and mangrove growth. To address this particularity, we utilized saltmarsh occurrence as absence data for mangroves (Cavanaugh et al., 2024; Enes Gramoso et al., 2026; Raw et al., 2023), as they both occur in similar geomorphological niches, being separated mainly by climate (Friess et al., 2012).

The package *biomod2* allows users to run different modeling algorithms simultaneously, compare their outputs and performance, and construct ensemble models. For this research, we used 10 different algorithms to account for variability in results. They were: artificial neural networks (ANN), classification tree analysis (CTA), flexible discriminant analysis (FDA), generalized additive models (GAM), generalized boosted models (GBM), generalized linear models (GLM), multivariate adaptive regression splines (MARS), random forest (RF), surface range envelope (SRE) and extreme gradient boosting (XBOOST). We split our data into random calibration (70% of data) and subsets for model cross-validation (30% of data). Models were built individually using these split subsets and repeated 30 times per algorithm, for a total of 100 runs for the ten algorithms. To access model performances, we used their total sum of squares (TSS), receiver operating characteristic (ROC) curves and Kappa statistics. Only models with TSS above 0.8 (TSS > 0.08) were retained to build the final ensemble model. We first made predictions of the current distribution of mangroves using current climate data, and then made predictions for future distribution of mangrove forests using future predicted data. Then, we compared the current and future predictions to assess changes in potential distribution. Binary predictions were obtained using the threshold that maximized TSS, as implemented in *biomod2*. After model prediction binarization, we compared current and future predictions and organized results into four categories: (I) mangrove stands predicted to persist, (II) mangrove stands predicted to be lost, (III) non-mangrove areas predicted to become occupied by mangroves and (IV) non-mangrove areas predicted to remain unoccupied.

### 3. Mangrove dispersal and connectivity

To estimate mangrove connectivity, we simulated propagule dispersal using OpenDrift v.1.13 (Dagestad et al., 2018), a Python-based software package based on Lagrangian method to simulate particle transport for modeling the trajectories and fate of objects drifting in the ocean. This software allows determining particle buoyancy, maximum age and interaction with the coastline, which is useful for simulating propagules. We used the Ocean Surface Current Analyses Real-time (OSCAR) database to access ocean surface current velocities (Dohan, 2021). OSCAR ocean velocities are calculated from satellite-sensed data. Daily averaged surface currents are available at a 0.25 x 0.25 degree grid from 1993 to present day.

As release points, we used the projected points (5km into the ocean). To access potential connectivity, we released 10^5^ particles from each release point. We assume these simulations to be an exaggeration of dispersal and propagule exchange between forests, as long distance dispersal (LDD) is estimated to account for only 1% - 5% of dispersal events (Van Der Stocken et al., 2019). Particles in the simulation had a maximum floating period of 120 days, which is a conservative estimate of the maximum viability observed in different mangrove propagules (Van Der Stocken et al., 2019). As propagules usually float passively on seawater surface with no information about sinking propagules for Western Atlantic mangrove species (Van Der Stocken et al., 2019), vertical mixing was disabled in the simulations. We used the snapped points (mangroves and saltmarshes) as stranding sites. We filtered stranded propagules using a 1,000 m buffer around each stranding site. We counted the stranded propagules inside each buffer and grouped results in a connectivity matrix. Finally, we binned propagule trajectories in a 10km X 10km grid to generate a map of the most frequent dispersal trajectories.

Using the obtained connectivity matrix from the oceanographic simulation (disconsidering selfloops), we estimated the degrees (in and out) and centrality (closeness and betweenness) of each sample point using the *igraph* v.2.1.1 package (Csárdi et al., 2026). For module detection, we used the *cluster_infomap* function (Rosvall & Bergstrom, 2008), as it takes into consideration the direction and weight of links, representatives of the source, destination and quantity of propagules. To do this, we excluded sites that did not release propagules (i.e. saltmarsh sites) and that did not receive propagules as they were, effectively, not part of the network and keeping them would affect network estimates. We also attributed modules to the nodes following the corresponding coastal sectors as classified by Schaeffer-Novelli (1990), which was recently reviewed (Cintrón-Molero et al., 2023). This allowed us to directly compare the algorithmic detection of modules (from *cluster_infomap*) with a climatological, hydrographic and oceanographic explicit sectorization (from the coastal classification).

To evaluate our simulated connectivity network, we generated 9,999 random networks by randomly permuting the connections (and keeping the network density constant) between mangroves in the observed connectivity matrix, according to the Erdõs-Rényi model. We compared the number of modules and modularity of the simulated and random networks using the same *cluster_infomap* function to detect the modules (Rosvall & Bergstrom, 2008). From this, we attributed the following categories to each sample point: (I) its module detected by the algorithm in the simulated connectivity, (II) its module in each of the 9999 random networks and (III) its module according to the coastal segment it belongs to. All points south of 32°35” (i.e., south of the segment VIII) were classified as segment IX. To measure how similar the modules detected in the networks and the coastal segments were in terms of how they group the same sites, we calculated the Adjusted Rand Index (ARI), using the *mclust* v.6.1.2 package (Scrucca et al., 2023). A value of 1 indicates identical partition between sets, 0.2 to 0.4 can be interpreted as moderate similarity in classification, whereas 0 indicates agreement expected by random chance. Using the coast segments as reference, an ARI value was calculated for the observed and and each of the random networks.

## Results

We observed based on the MapBiomas dataset that the southernmost mangrove forest in the Atlantic coast of South America occurs at 28°28’68” S (Fig. 1). South to that limit, the Brazilian coast consists mostly of sandy beaches, coastal lagoons and rocky shores, which are usually unsuitable for propagule establishment, until the Lagoa dos Patos estuary, at around 31°S. Overall, expectedly, minimum sea surface temperature (SST) decreases as latitude increases, but between approximately 20°S and 30°S, SST is colder in some points, possibly because of the Cabo Frio upwelling in this region (Valentin, 1984). South to 30°S, SST drops drastically, likely due to the Malvinas Current, which flows northward from the pole, bringing cold water and air masses (Maamaatuaiahutapu et al., 1998; Piola & Matano, 2019).

### 1. Environmental niche modeling

Algorithms from our environmental niche model performed well (TSS > 0.835; ROC > 0.918; Kappa > 0.784; Table S1), thus, we retained all but one algorithm (RF was excluded to avoid overfitting; see Table S1) to build the ensemble model. The ensemble model closely matched the current distribution of mangrove stands in South America (Fig. 2), with high probability of mangrove habitat suitability where they currently occur and near zero probability where they are absent (i.e saltmarsh areas). The threshold for model binarization was set at 651, as that is the value that maximized TSS. For current predictions, two sites unoccupied nowadays by mangroves but within their present distribution were detected as suitable (Points 144 and 146).

**Figure 2.**
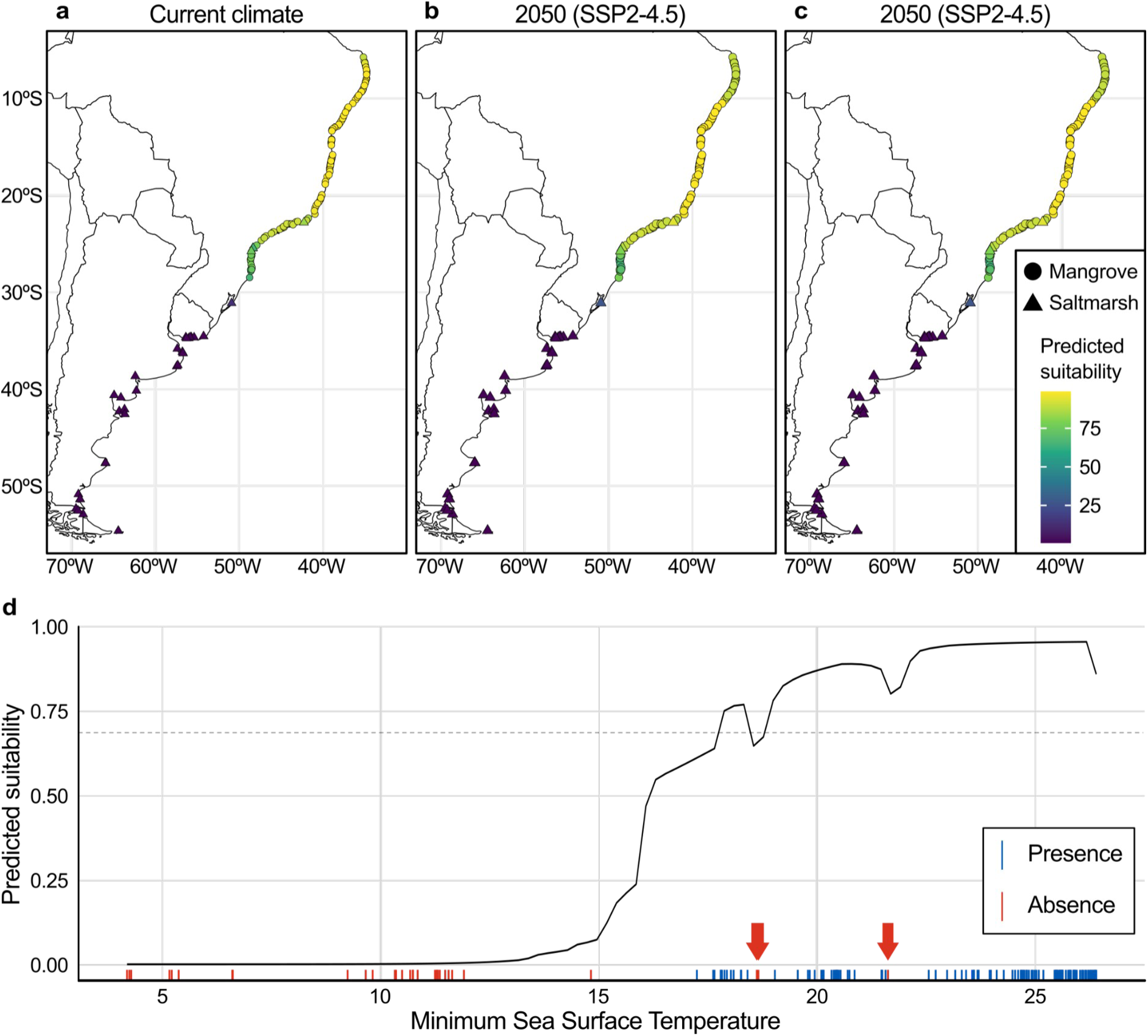
Environmental suitability of Atlantic South American mangrove and saltmarsh sites under current climate (a) and future climate scenarios (b: SSP2-4.5; c: SSP5-8.5). Response curves for ensemble (d) models, with the gray dashed line indicating binarization threshold (651). Note arrows indicating absence points that generated low predicted suitability valleys.

Future predictions from our environmental niche models were similar between different climate change scenarios. In both SSP2-4.5 and SSP5-8.5, increase in habitat suitability that indicated mangrove gain was predicted only within the current mangrove range (Figs. 2 and 3). Contrary to our expectations, habitat suitability was not predicted to increase substantially beyond the current distribution of mangroves, and no sites southward to its current distribution were deemed suitable for mangrove development by 2050. The model also indicated decreased suitability at multiple mangrove sites in both climate scenarios (Fig. 2) and, after binarization, mangrove loss at these estuaries (Fig. 3).

**Figure 3.**
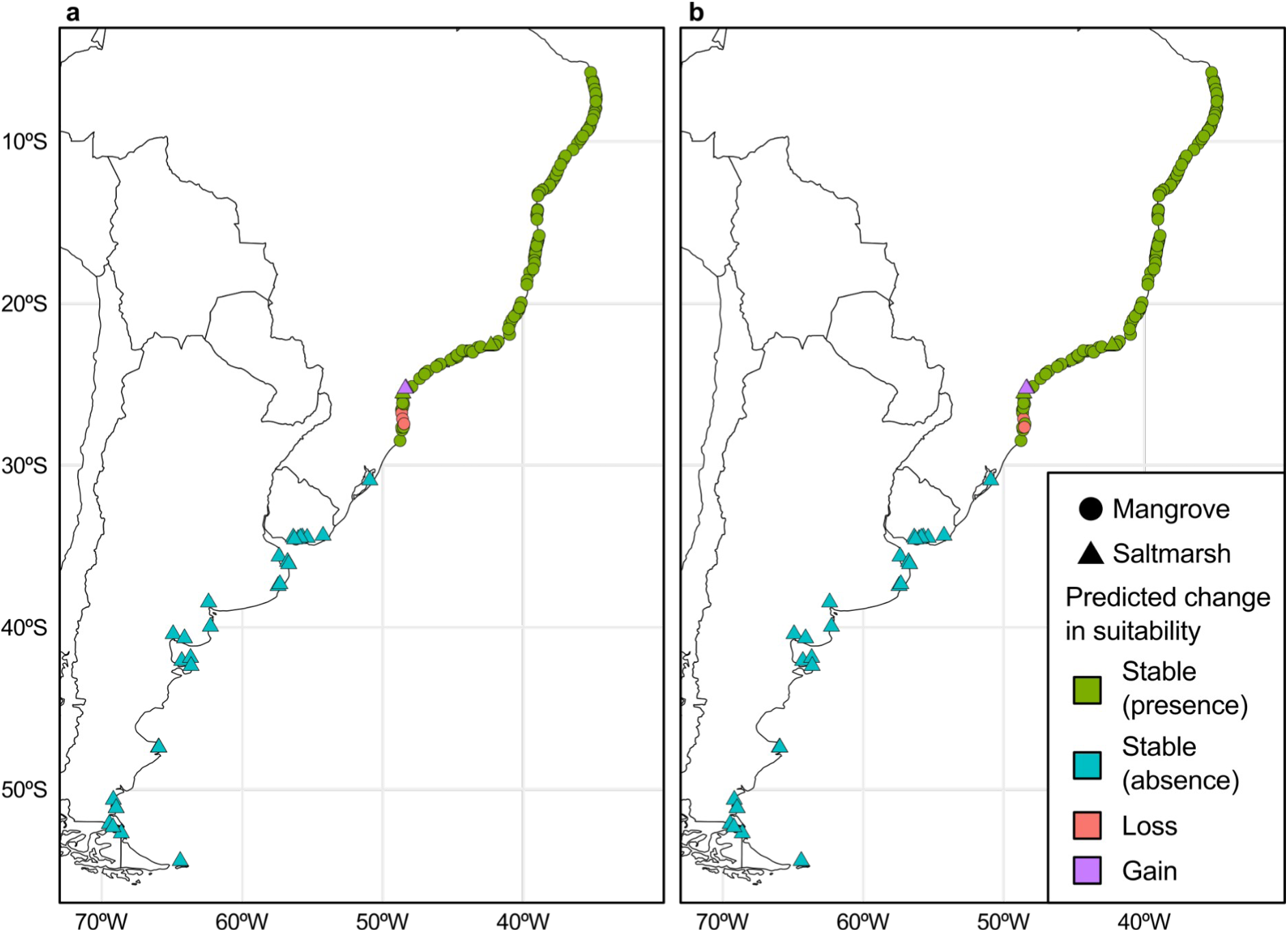
Binary predictions for the occurrence of Atlantic mangroves of South America under SPP2-4.5 (a) and SSP5-8.5 (b) future climate scenarios.

### 2. Mangrove dispersal and connectivity

Generally, we observed that propagule dispersal direction occurred southwards and stays relatively close to the continent, and propagule *in situ* retention is frequent (Fig. 4; as indicated by the high density diagonal in Fig. 5 and Table S3). Despite these overall findings, direction of dispersal showed a wide variability (colored spots above the high density diagonal of the matrix indicate northward dispersal, whereas points below the diagonal indicate southwards dispersal), with some regions (around 5°S to 10°S) dispersing mostly northwards and others (around 10°S to 20°S) dispersing frequently in both directions. Beyond the current range limit of mangroves (28°28’68” S), our results showed that only three sites served as stranding areas for propagules. The first is Lagoa dos Patos (31°15’35” S; Point 147), the largest choked coastal lagoon in the world (Kjerfve, 1986). Even though a sandbar separates Lagoa dos Patos from the Atlantic Ocean, we found that propagules from multiple sources would be able to enter the lagoon through the Rio Grande inlet (Fig 4). Beyond Lagoa dos Patos, few propagules reach the coastline and stranding becomes rare, due to the Malvinas Current, which branches from the Antarctic Circumpolar Current that circulates around the south pole, and flows northward along the South American coast. Despite this counter-directional flow, we found that propagules also could strand at two other sites: (I) near the Rio da Prata estuary (37°47’7” S; Point 123) and (II) in the Strait of Magellan (52°28’8” S; Point 148), at the southernmost tip of South America (Fig. 4). Despite the potential for propagules to strand in these areas, these should be considered rare events, as few (< 3000 stranded particles out of the 1.2M released) propagules reached this far over ten years of simulation.

**Figure 4.**
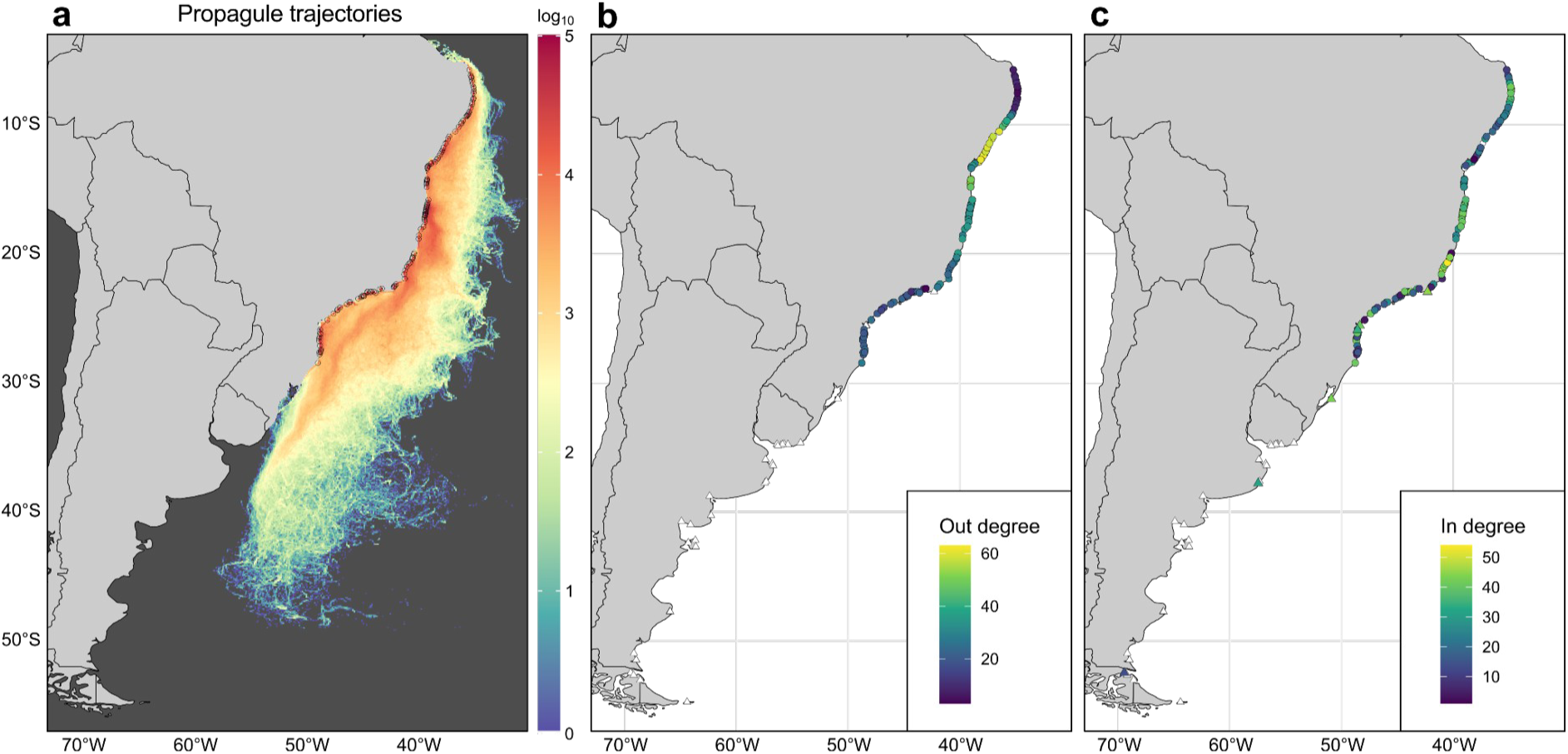
Simulated trajectory density map of mangrove propagules released along the Atlantic coast of South America (a). 105 propagules were released from each mangrove site (blue points) over a ten year simulation period (2010-2020). Propagule trajectories were binned in a 10 km x 10 km grid. Map of degrees in (b) and out (c) of mangrove and saltmarsh sites.

**Figure 5.**
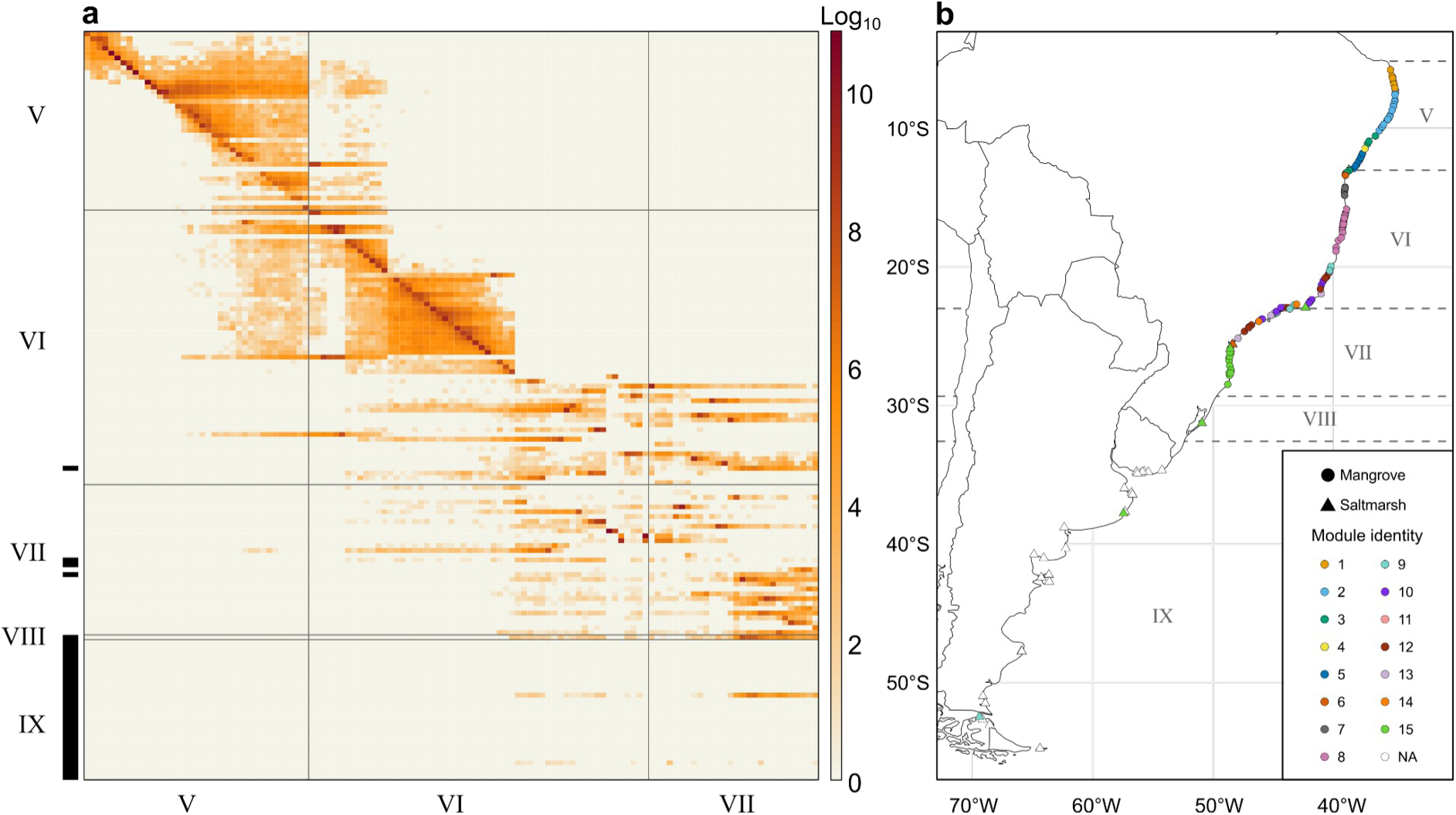
Connectivity matrix between present mangrove sites (x-axis) and stranding locations (y-axis) (a). Coast sectors are represented by roman numerals. Black rectangles indicate saltmarsh sites. Modules of mangrove and saltmarsh sites as detected in our network analyses (b), compared to coastal segments (Schaeffer-Novelli et al., 1990).

From our network analyses, we found that mangrove forests varied greatly in their connectivity (Figs. 4 and 5). Most sites received more propagules than they retained (89 sites), whereas 34 sites retained more propagules than they received from other sources (Table S3). Not considering propagule retention (i.e. stranding at the origin site), the strongest disperser was at Estação Ecológica de Guanabara (22°41’38” S; Point 108), a conservation unit in Rio de Janeiro, which successfully dispersed 41,978 propagules to other mangroves, and Guarapari (20°41’11” S; Point 98) dispersed to 64 different sites. As for propagule stranding, Guarujá (23°55’25” S; Point 109) received the most propagules from other sites, at 45,227, but Itacaré (14°20’42” S; Point 31) received propagules from the most different mangrove forests, at 53 (Table S3). From our connectivity data, we detected 15 modules in the network, whereas random networks varied from 1 to 23 modules (mean = 16.27±2.26) (Fig. S2). Our observed network had high modularity (0.73), higher than expected by random networks (0.51±0.04) and coastal segments alone (0.25) (Fig. S3). Comparison between the inferred modules and the coast segments shows an agreement significantly higher than expected by chance (ARI = 0.25; mean ARI of random networks = - 9.03 x 10^-5^; p = 10^-4^; Table S3).

## Discussion

We investigated the potential for shifts in the distribution of South West Atlantic mangroves in different climate change scenarios using complementary modelling techniques. Overall, our results did not support the hypothesis that increased temperatures due to climate change would lead to the poleward expansion of South American mangrove forests. We found that minimum SST acted as a major driver of the range limit mangroves and that it will continue to act by 2050 (under both SSP2-4.5 and SSP5-8.5), limiting expansion beyond the current observed distribution. Dispersal acted as a secondary barrier, heavily limiting propagule stranding beyond 35°S.

From our habitat suitability predictions, we found that some non-mangrove sites within the current mangrove range are already suitable for mangrove development or could become so by 2050. These estuaries are currently occupied by saltmarshes, indicating that mangrove encroachment into these sites may depend on the interactions between mangroves and saltmarshes. Saltmarsh vegetation facilitates mangrove propagule recruitment, but may also slow seedling and tree growth through competition (Hockaday et al., 2023; Reis et al., 2023, 2025). Nevertheless, mangrove encroachment into saltmarsh-dominated areas is well reported (Coldren & Proffitt, 2017; Howard et al., 2015), and, excluding sites with low temperatures, mangroves seem to be superior competitors (Reis et al., 2023). We found that these areas showed high potential of receiving propagules from different mangrove stands, however, our simulations represent an extrapolation of long distance dispersal, which is relatively uncommon in nature, accounting for only 1 - 5% of dispersal events (Van Der Stocken et al., 2019). Consequently, the realized influx of propagules in these sites may have been insufficient to surpass the competitive pressure exerted by saltmarshes after early seedling establishment (Reis et al., 2023), preventing the development of full mangrove stands, an ecological question that deserved further investigation.

We found no sites beyond the present distribution of mangroves that are suitable for mangrove development, neither under contemporary nor future (SSP2-4.5 and SSP5-8.5) minimum sea surface temperatures. Mangrove physiology may be limited under conditions like low air or water temperatures and hypersalinity, but different species have different tolerances. At the southernmost South-American latitudinal limit, *Rhizophora mangle* stops occurring at a lower latitude (27°38’ S) than *Laguncularia racemosa* and *Avicennia schaueriana* (28°28’ S) (Soares et al., 2012). *R. mangle* propagules have longer floating periods, but *L. racemosa* seems to be more cold-tolerant (Saintilan et al., 2014), which could explain the difference in range between species. Still, these mangrove trees at their range edge present stunted growth and development (Soares et al., 2012), possibly as a reflection of the physiological limits of these species. The Atlantic coast of South America has a clear temperature gradient that becomes more steep around the range limit of mangroves (25° S - 30° S), with SST dropping drastically beyond 30° S (Fig. 1) and chilling events (where temperatures drop below 15°C) becoming much more frequent starting at around 29° S (A. C. Ximenes et al., 2021). These climatic conditions may be explained by the Brazil-Malvinas confluence zone, and our results suggest that cold temperatures are likely the main limiting factor for the distribution of mangrove forests in South America.

Unexpectedly, our model predicted loss of habitat suitability in some mangrove sites near their southern range edge. This result appeared in both future climate scenarios, but sites of predicted loss varied slightly in latitude between SSP2-4.5 and SSP5-8.5. High temperatures and mangrove dieback have only been associated under arid conditions (Adame et al., 2021), and little is known about an upper boundary of temperature tolerance for mangrove trees. This does not seem to be the case for our predictions, as mangroves in these southern sites are historically exposed to SST minima below 20°C (around 18°C), and future scenarios predict an increase in minimum SST ranging from 0.7°C (SSP2-4.5) to 0.8°C (SSP5-8.5) (Fig. 1), which would remain within ranges of mangrove thermal tolerance. After careful examination, we suggest that these results could be an artifact of our model, which estimates probability of occurrence from presence and absence data along a temperature gradient. Inspection of our presence and absence dataset reveals two non-mangrove sites that occur at around 19°C (Fig. 2C and 2D, arrow). Because response curves vary across algorithms, from simple sigmoidal shapes to more complex and non-monotonic forms, local valleys of low predicted probability may emerge due to absence points. When building the ensemble model, these valleys were retained. As a result, sites that are currently predicted to have moderate to high probabilities of mangrove occurrence (at around 18°C) fell into a valley of low probability when shifted to slightly higher temperatures (at around 19°C), leading to predictions of mangrove loss that do not necessarily correspond to the physiological limits of mangrove trees.

We observed the potential for dispersal beyond the current range of mangroves in our simulations, as propagules reached three currently unoccupied sites: Lagoa dos Patos (31°15’35” S; Point 147), near Rio da Prata (37°47’7” S; Point 123) and the Strait of Magellan (52°28’8” S; Point 148) (Fig. 5; Table S3). The latter two occur at elevated latitudes, where propagule stranding is infrequent and SST remains well below suitable conditions, making mangrove encroachment unlikely in these areas. Although suitability increased slightly at Lagoa dos Patos by 2050, this was still insufficient for it to be considered a viable site for mangrove development. However, if low temperature or chilling event frequency alleviate further than expected, Lagoa dos Patos would likely represent the most probable site for mangrove latitudinal expansion in South America. Under such conditions, more cold-tolerant species, such as *L. racemosa* (Santos Borges et al., 2019), might have a greater chance at expanding their range.

Although propagules in our simulation were able to reach maximum latitudes of around 50° S, stranding became very rare beyond 35° S. This result may reflect the Brazil-Malvinas confluence, where the southward-flowing Brazil current meets the northward Malvinas current around 35° S - 40° S (varying seasonally) (Piola & Matano, 2019). At this region, the Brazil current is mainly redirected about 45° east of its original poleward trajectory (Maamaatuaiahutapu et al., 1998; Piola & Matano, 2019). The Brazil-Malvinas confluence zone not only generates steep temperature gradients (Fig. 1) (Piola & Matano, 2019), but also creates a major barrier for dispersal, by redirecting propagules to the middle of the Atlantic ocean (Fig. 4). Future changes in ocean circulation could potentially modify this barrier. Some evidence suggests an intensification of the South Atlantic Gyre (Taschetto & Wainer, 2003), which feeds the Brazil current, and a southward migration of the Antarctic Circumpolar Current (Biastoch et al., 2009), which could affect the Brazil-Malvinas confluence zone. With the intensification of these trends in the long term, it is possible that propagule stranding becomes more frequent in higher latitudes. Still, the consequences for potential range shifts remain uncertain, as geomorphologically and climatically suitable estuaries are infrequent beyond the current southernmost distribution limit in Laguna and propagules reaching these regions would likely face long floating periods and low SST, conditions that would compromise their viability.

We found that propagule retention is high, but connectivity between mangrove forests is also frequent. Based on dispersal connections, our sampling sites can be divided into 15 modules, indicating that propagule exchange may happen more frequently within certain groups than along the entire coast. The agreement we found between the modules and the coast segments (Cintrón-Molero et al., 2023; Schaeffer-Novelli et al., 1990), support the idea that these segments have, at least partially, effects on dispersal processes. Despite these modular and spatial structures, *R. mangle* (Madeira et al., 2023)(Francisco 2018), *A. schaueriana* (Cruz et al., 2019; Mori et al., 2015) and *L. racemosa* (Sereneski-Lima et al., 2021) individuals within the study area appear to form a somewhat genetically homogenous population. For *R. mangle,* directional gene flow estimates were performed and, although no evidence for recent gene flow has been found, there was evidence of long-term migration rates (Madeira et al., 2023). Possibly, the modular structure we observed is not strong enough for genetic differences to accumulate between groups, but, instead, helps promote genetic mixing along the heterogeneous Brazilian coast.

Overall, we found no evidence of current nor future poleward expansion of mangroves, but total mangrove area has increased in Brazil over the last decades (Vanin et al., 2025b), likely due to inland expansion, and dynamic coastal processes. Landward migration of mangroves is expected as a response to rising sea levels (Godoy & Lacerda, 2015), but anthropogenic interference and urban expansion may restrict this process, leading to coastal squeeze (Almeida et al., 2025). Under scenarios where both landward and poleward migration are limited, South American mangroves may face increasing vulnerability, with potential impacts on the several ecological, biogeochemical and social (Barbier et al., 2011; Lee et al., 2014) cycles they support.

## Conclusion

Low temperatures seem to be the main factor limiting the poleward expansion of mangroves in the Atlantic coast of South-America and there is evidence other limiting factors will not emerge by 2050. Our results indicate that minimum SST strongly constrains habitat suitability beyond the current distribution of mangroves, while dispersal processes act as a secondary barrier by limiting propagule stranding beyond approximately 35° S. In particular, the Brazil–Malvinas confluence creates both steep temperature gradients and an oceanographic barrier that restricts propagule stranding in higher latitudes. Also, geomorphologically suitable sites are much more sparse beyond the current range limit, which limits stranding viability. Together, these climatic, oceanographic and geomorphological factors help explain why the poleward expansion reported in other regions(Friess et al., 2022; Hickey et al., 2017) has not yet been observed in the South American range limit of mangrove forests.

The abrupt limit of mangroves in the Atlantic coast of South America has received the attention of multiple researchers over the years. Our results contribute to leading hypotheses of climate restriction and shed light on the role of ocean currents. Still, future research should investigate other possible drivers of the observed range limit (see Molero & Novelli, 2019) and how they might act in tandem with human-driven changes in mangrove distribution.

## Acknowledgments

We thank Camila Castanho and Silas Poloni for discussions and comments on the development of this research project.

## Author Contributions

Miguel P. Pereira-Romeiro: Conceptualization (lead); Data curation (lead); Formal analysis (lead); Funding acquisition (lead); Investigation (lead); Methodology (lead); Project administration (lead); Software (lead); Validation (equal); Visualization (lead); Writing – original draft (lead); Writing – review and editing (equal). Gustavo M. Mori: Conceptualization (equal); Funding acquisition (supporting); Investigation (equal); Methodology (equal); Supervision (equal); Validation (equal); Writing – review and editing (equal). Flávia M. D. Marquitti: Conceptualization (equal); Formal analysis (supporting); Funding acquisition (equal); Investigation (equal); Methodology (equal); Project administration (equal); Software (supporting); Supervision (lead); Validation (equal); Writing – review and editing (equal). All authors provided editorial advice, contributed critically to drafts and gave final approval for publication

