## Supplementary Material for "Range shifts of Eastern South American mangroves in a changing climate"

### Supplementary information

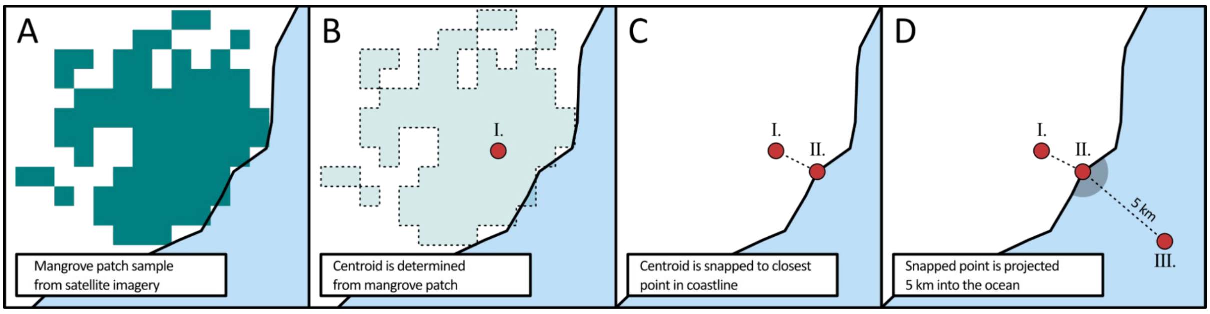


**Figure S1.** Schematic showing transformations made to sample points. Centroids (I) were used for ecological niche models. Projected points (III) were used as release points for propagule dispersion simulations. Snapped points (II) were used as stranding sites.

**Table S1.** Algorithms mean performance (± standard deviation). Each algorithm was run 30 times. Algorithms are: artificial neural networks (ANN), classification tree analysis (CTA), flexible discriminant analysis (FDA), generalized additive models (GAM), generalized boosted models (GBM), generalized linear models (GLM), multivariate adaptive regression splines (MARS), random forest (RF), surface range envelope (SRE) and extreme gradient boosting (XBOOST). RF was not retained to build the ensemble model, as its performance is likely an artifact of overfitting.

| Algorithm | Kappa | ROC | TSS |
|---|---|---|---|
| ANN | 0.938 (0.025) | 0.982 (0.008) | 0.906 (0.037) |
| CTA | 0.934 (0.033) | 0.964 (0.018) | 0.919 (0.032) |
| FDA | 0.938 (0.025) | 0.953 (0.019) | 0.906 (0.037) |
| GAM | 0.938 (0.025) | 0.982 (0.008) | 0.906 (0.037) |
| GBM | 0.996 (0.010) | 1.000 (0.000) | 0.995 (0.013) |
| GLM | 0.938 (0.025) | 0.982 (0.008) | 0.906 (0.037) |
| MARS | 0.969 (0.023) | 0.998 (0.002) | 0.976 (0.021) |
| RF | 1.000 | 1.000 | 1.000 |
| SRE | 0.784 (0.026) | 0.918 (0.019) | 0.835 (0.037) |
| XGBOOST | 0.980 (0.017) | 0.999 (0.002) | 0.976 (0.020) |

**Table S2.** Evaluation metric for the ensemble model. TSS cutoff was used for prediction binarization.

| Evaluation metric | Cutoff | Sensitivity | Specificity | Calibration |
|---|---|---|---|---|
| TSS | 651.0 | 100 | 97.774 | 0.968 |
| ROC | 648.5 | 100 | 97.774 | 0.996 |
| Kappa | 651.0 | 100 | 97.774 | 0.979 |

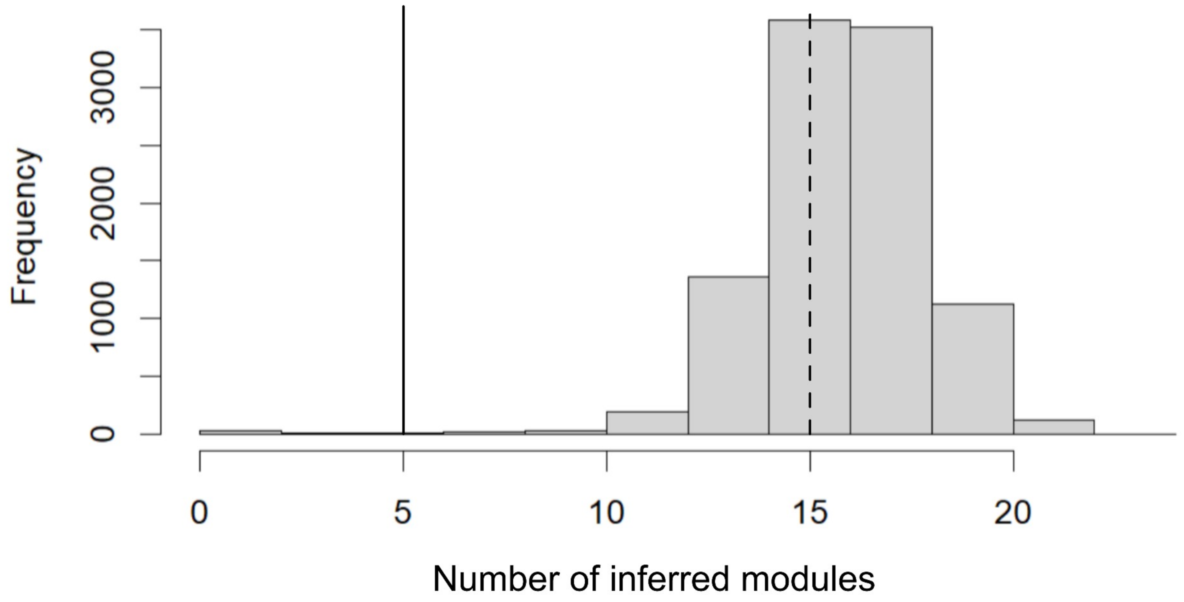


**Figure S2.** Histogram of number of inferred modules of random networks (N=9999). The continuous line indicates the number of modules defined by coastal segments alone. The dashed line indicates the number of modules inferred from the observed network obtained from simulations.

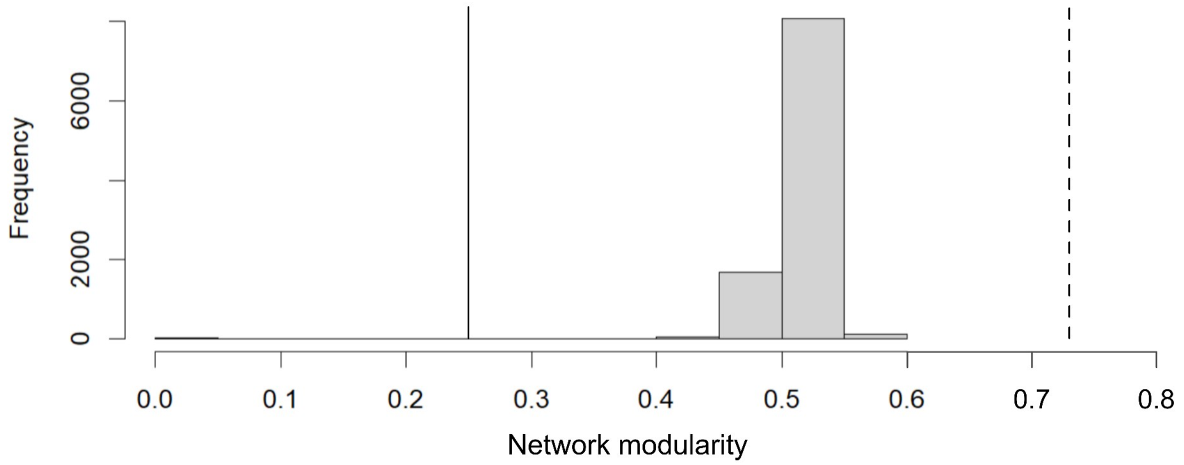


**Figure S3.** Histogram of inferred modularity of random networks (N=9999). The continuous line indicates the modularity inferred from coastal segments alone (0.25). The dashed line indicates the modularity inferred from the observed network obtained from simulations (0.73).

**Table S3.** Connectivity overview of Atlantic mangrove and saltmarsh sites in South America. “Type” column indicates vegetation type, with M indicating mangrove and S indicating saltmarsh. “Disp. prop.” stands for propagules that were successfully dispersed to other sites. “Strand. prop.” represents propagules stranded in that site. “Prop. retain.” shows propagules that originated and stranded in that same site. “Mod.” indicates the community identity found through our modularity analyses. “Out deg.” and “In deg.” represent out degree and in degree of each site, respectively.

| ID | Type | Longitude | Latitude | Sector | Disp. prop. | Strand. prop. | Prop. retain. | Mod. | Out deg. | In deg. |
|---|---|---|---|---|---|---|---|---|---|---|
| 1 | M | -35.3869117023937 | -9.2859301465170 | 1 | 10193 | 3525 | 1032 | 2 | 22 | 22 |
| 2 | M | -35.0343309835010 | -6.2834251696750 | 1 | 4528 | 1928 | 871 | 1 | 9 | 19 |
| 3 | M | -35.9602951543722 | -9.9371249024557 | 1 | 4676 | 1526 | 1410 | 2 | 41 | 24 |
| 4 | M | -34.8287218404382 | -7.9546857406084 | 1 | 242 | 7409 | 46267 | 2 | 4 | 40 |
| 5 | M | -35.3436917483953 | -9.2349123405216 | 1 | 7568 | 7703 | 1017 | 2 | 20 | 23 |
| 6 | M | -34.9656430924889 | -6.5967944856645 | 1 | 655 | 1553 | 46822 | 1 | 8 | 24 |
| 7 | M | -35.2249160014023 | -9.0240127335379 | 1 | 4325 | 3790 | 1547 | 2 | 15 | 26 |
| 8 | M | -34.8581582504359 | -8.0264230636037 | 1 | 984 | 15761 | 7700 | 2 | 7 | 38 |
| 9 | M | -36.1514866123644 | -10.1578082504346 | 1 | 2966 | 525 | 2599 | 2 | 44 | 16 |
| 10 | M | -35.4892227763909 | -9.3780140555076 | 1 | 7040 | 2596 | 1073 | 2 | 28 | 23 |
| 11 | M | -35.2940696043978 | -9.1589827885278 | 1 | 5096 | 4645 | 812 | 2 | 21 | 23 |
| 12 | M | -35.4283381213926 | -9.3216590565133 | 1 | 5615 | 4129 | 6538 | 2 | 29 | 23 |
| 13 | M | -34.8016430814651 | -7.2064062596443 | 1 | 1244 | 1292 | 24584 | 1 | 8 | 40 |
| 14 | M | -34.8048501914580 | -7.4029732596353 | 1 | 478 | 2365 | 53151 | 2 | 6 | 42 |
| 15 | M | -35.9016141333748 | -9.8621494494626 | 1 | 6372 | 141 | 388 | 2 | 42 | 13 |
| 16 | M | -34.9393737194286 | -8.2483660735895 | 1 | 1027 | 22762 | 12574 | 2 | 8 | 37 |
| 17 | M | -35.7683439493807 | -9.6903555794781 | 1 | 4198 | 3815 | 1565 | 2 | 39 | 21 |
| 18 | M | -35.0946609335053 | -6.1819701526768 | 1 | 1163 | 4647 | 2062 | 1 | 9 | 13 |
| 19 | M | -35.2024485425220 | -5.7509902966902 | 1 | 379 | 983 | 1803 | 1 | 7 | 10 |
| 20 | M | -34.9688417794229 | -8.4102832525802 | 1 | 10082 | 4854 | 1702 | 2 | 9 | 31 |
| 21 | M | -35.0827695554140 | -8.6794672925619 | 1 | 3577 | 404 | 99 | 2 | 8 | 21 |
| 22 | M | -35.0262927424983 | -6.3552669546723 | 1 | 3619 | 1336 | 12144 | 1 | 9 | 19 |
| 23 | M | -34.9150299054812 | -6.7947135096579 | 1 | 765 | 1489 | 48460 | 1 | 10 | 32 |
| 24 | M | -34.8410568754707 | -7.0629785066492 | 1 | 324 | 2475 | 12664 | 1 | 6 | 42 |
| 25 | M | -34.8301048324530 | -7.5504370966274 | 1 | 188 | 2040 | 556 | 2 | 4 | 40 |
| 26 | M | -39.1742402730537 | -17.0530652798178 | 2 | 7868 | 6973 | 8257 | 8 | 30 | 41 |
| 27 | M | -39.1241487066151 | -16.7282175775027 | 2 | 6137 | 13338 | 4283 | 8 | 33 | 42 |
| 28 | M | -39.1547934690582 | -16.9642906308257 | 2 | 6852 | 8334 | 10145 | 8 | 27 | 43 |
| 29 | M | -38.0313004962642 | -12.5974860431955 | 1 | 3414 | 1779 | 946 | 5 | 60 | 28 |
| 30 | M | -38.9651881922281 | -13.2942289071015 | 2 | 12524 | 6034 | 12355 | 6 | 26 | 32 |
| 31 | M | -39.0008518629554 | -14.3451055923105 | 2 | 3555 | 1394 | 4175 | 7 | 55 | 27 |
| 32 | M | -39.0349582751728 | -14.5125091010135 | 2 | 2785 | 2452 | 10959 | 7 | 52 | 32 |
| 33 | M | -39.0247485230873 | -16.3802746148774 | 2 | 6012 | 8918 | 3249 | 8 | 33 | 44 |

| | | | | | | | | | | |
|---|---|---|---|---|---|---|---|---|---|---|
| 34 | M | -39.2134651800476 | -17.1697396438067 | 2 | 4165 | 5787 | 11326 | 8 | 32 | 39 |
| 35 | M | -39.0503771821695 | -14.5838929290076 | 2 | 3092 | 1695 | 4817 | 7 | 50 | 32 |
| 36 | M | -39.0865433210782 | -16.5573277458606 | 2 | 6415 | 3436 | 6067 | 8 | 30 | 37 |
| 37 | M | -38.9881920241867 | -14.2173280570368 | 2 | 2793 | 2980 | 3124 | 7 | 60 | 26 |
| 38 | M | -39.0182259051774 | -14.4192532670211 | 2 | 2870 | 1175 | 4697 | 7 | 59 | 26 |
| 39 | M | -39.1048610650719 | -16.6863243868498 | 2 | 5757 | 7590 | 9534 | 8 | 27 | 42 |
| 40 | M | -38.2727286392510 | -12.8760093661650 | 1 | 2694 | 1390 | 3584 | 5 | 58 | 30 |
| 41 | M | -39.1712965840558 | -17.0091021478213 | 2 | 6784 | 9980 | 5756 | 8 | 31 | 42 |
| 42 | M | -39.1453035530656 | -16.8103553748381 | 2 | 7561 | 8552 | 11326 | 8 | 32 | 45 |
| 43 | M | -38.9939314975372 | -14.2698058132969 | 2 | 3437 | 2824 | 3004 | 7 | 50 | 26 |
| 44 | M | -38.9443695541022 | -16.0755127069046 | 2 | 3849 | 4862 | 5121 | 8 | 31 | 35 |
| 45 | M | -37.6195506132921 | -11.9717597452560 | 1 | 2351 | 2142 | 724 | 5 | 63 | 26 |
| 46 | M | -37.7666783712807 | -12.2324130832322 | 1 | 4068 | 745 | 136 | 5 | 59 | 16 |
| 47 | M | -38.9723923380963 | -16.1995893018937 | 2 | 5474 | 2831 | 2523 | 8 | 36 | 37 |
| 48 | M | -38.9339421372315 | -13.2227101601079 | 2 | 4110 | 19616 | 6662 | 6 | 19 | 34 |
| 49 | M | -37.8780512912738 | -12.3875460622167 | 1 | 4269 | 63 | 38 | 5 | 61 | 9 |
| 50 | M | -37.6800865422872 | -12.0862790252458 | 1 | 2673 | 3833 | 1247 | 5 | 58 | 24 |
| 51 | M | -39.0203438040924 | -16.2691175428859 | 2 | 4924 | 3405 | 3932 | 8 | 30 | 39 |
| 52 | M | -39.1275375490622 | -16.8872126548331 | 2 | 7849 | 4394 | 1536 | 8 | 33 | 42 |
| 53 | M | -39.2139396260389 | -17.3531527107923 | 2 | 3311 | 5676 | 10237 | 8 | 33 | 40 |
| 54 | M | -38.1162100202594 | -12.7003328681845 | 1 | 4347 | 1 | 1 | 5 | 61 | 1 |
| 55 | M | -39.5458824319986 | -18.0874815017161 | 2 | 1808 | 570 | 2352 | 8 | 34 | 27 |
| 56 | M | -39.0628185330835 | -16.4523317628699 | 2 | 7829 | 4067 | 2354 | 8 | 32 | 35 |
| 57 | M | -39.1920025200309 | -17.5259396707800 | 2 | 2921 | 4645 | 11860 | 8 | 33 | 41 |
| 58 | M | -39.7296683449715 | -18.5791099986663 | 2 | 1552 | 264 | 2507 | 8 | 28 | 24 |
| 59 | M | -39.0246790031588 | -14.8224862389921 | 2 | 2457 | 629 | 8701 | 7 | 47 | 27 |
| 60 | M | -39.7484089069580 | -18.8448620086436 | 2 | 1124 | 860 | 4091 | 8 | 36 | 27 |
| 61 | M | -37.5331383453002 | -11.7825731222721 | 1 | 2041 | 1131 | 946 | 5 | 60 | 16 |
| 62 | M | -36.4677259463509 | -10.5287917903987 | 1 | 2559 | 1138 | 5256 | 3 | 60 | 19 |
| 63 | M | -38.6436568782427 | -13.0152960671368 | 2 | 6538 | 7258 | 5348 | 3 | 25 | 29 |
| 65 | M | -38.6436568782427 | -13.0152960671368 | 2 | 11451 | 0 | 0 | 3 | 29 | NA |
| 66 | M | -37.1235564933289 | -11.0969161223339 | 1 | 4175 | 12591 | 15 | 3 | 59 | 29 |
| 67 | M | -37.0037951233362 | -10.9146142383506 | 1 | 4593 | 1724 | 475 | 3 | 57 | 18 |
| 68 | M | -38.8549590542344 | -13.1705321411155 | 2 | 9264 | 2210 | 0 | 3 | 35 | 14 |
| 69 | M | -39.3081785800120 | -17.8852972977452 | 2 | 5744 | 15596 | 0 | 8 | 35 | 40 |
| 70 | M | -38.8688835101151 | -15.8184900659278 | 2 | 3181 | 5696 | 13 | 8 | 37 | 37 |
| 71 | M | -37.3413389463142 | -11.4518616433019 | 1 | 1891 | 0 | 0 | 4 | 58 | NA |
| 72 | M | -38.9588521862247 | -13.3704345630966 | 2 | 13227 | 0 | 0 | 6 | 29 | NA |
| 73 | M | -41.8857983607171 | -22.4859126772095 | 2 | 3578 | 4770 | 0 | 6 | 29 | 32 |
| 74 | M | -44.0007103476243 | -22.9423601220452 | 2 | 4633 | 4401 | 15 | 11 | 28 | 43 |
| 75 | M | -40.3219264438669 | -20.4210790134797 | 2 | 3157 | 3007 | 0 | 11 | 34 | 14 |
| 76 | M | -44.0789620956208 | -22.9562778530393 | 2 | 3206 | 2950 | 0 | 11 | 28 | 25 |

| | | | | | | | | | | |
|---|---|---|---|---|---|---|---|---|---|---|
| 77 | M | -40.9766840377717 | -21.9305952133112 | 2 | 3057 | 317 | 0 | 13 | 28 | 6 |
| 78 | M | -44.8386040605681 | -23.3690772679569 | 3 | 20106 | 3184 | 0 | 13 | 18 | 30 |
| 79 | M | -44.8880414445680 | -23.3392500819567 | 3 | 868 | 20511 | 2296 | 13 | 13 | 33 |
| 80 | M | -45.1660939391353 | -23.4926004359261 | 3 | 946 | 876 | 183 | 13 | 22 | 17 |
| 81 | M | -47.9189087733190 | -25.1457166126097 | 3 | 3926 | 64 | 0 | 13 | 25 | 3 |
| 82 | M | -44.6367936355836 | -23.2378083229808 | 3 | 1261 | 2005 | 0 | 13 | 24 | 22 |
| 83 | M | -44.3097884236117 | -22.9703259660244 | 2 | 35209 | 88 | 0 | 12 | 6 | 7 |
| 84 | M | -46.9998100004169 | -24.3310509997399 | 3 | 6083 | 13819 | 6 | 10 | 15 | 21 |
| 85 | M | -45.9660036694974 | -23.7819413998521 | 3 | 6384 | 6944 | 17 | 10 | 22 | 44 |
| 86 | M | -40.9609119328069 | -21.2876311353686 | 2 | 13820 | 9289 | 2 | 10 | 25 | 48 |
| 87 | M | -40.8070722368258 | -21.0014636464022 | 2 | 3245 | 16166 | 351 | 10 | 21 | 43 |
| 88 | M | -41.9434272887131 | -22.5279656712025 | 2 | 16045 | 7101 | 0 | 10 | 28 | 39 |
| 89 | M | -41.7666163997276 | -22.3562520032281 | 2 | 1117 | 15269 | 3 | 10 | 30 | 26 |
| 90 | M | -40.2871931038731 | -20.3153482104906 | 2 | 2648 | 1404 | 10 | 10 | 31 | 19 |
| 91 | M | -40.1854682648893 | -20.0416132305195 | 2 | 1655 | 2003 | 41 | 9 | 33 | 12 |
| 92 | M | -44.6555941025795 | -23.2945512839746 | 3 | 1442 | 760 | 0 | 9 | 23 | 22 |
| 93 | M | -44.7149902665824 | -23.2104726639787 | 3 | 1214 | 7 | 0 | 10 | 19 | 4 |
| 94 | M | -41.9858827597088 | -22.5822247711952 | 2 | 13241 | 88 | 0 | 10 | 27 | 5 |
| 95 | M | -45.8827096015021 | -23.7622187038589 | 3 | 11839 | 12985 | 0 | 10 | 24 | 26 |
| 96 | M | -44.3335030006134 | -22.9278445220268 | 2 | 11402 | 12383 | 0 | 10 | 10 | 41 |
| 97 | M | -46.7911120004356 | -24.1875189997656 | 3 | 6698 | 34747 | 1 | 12 | 16 | 12 |
| 98 | M | -40.5074224378491 | -20.6864552804464 | 2 | 4112 | 9258 | 7 | 12 | 25 | 54 |
| 99 | M | -47.3743960003786 | -24.6480369996882 | 3 | 6857 | 4701 | 0 | 12 | 20 | 42 |
| 100 | M | -41.0471198887885 | -21.5902779273372 | 2 | 3017 | 7132 | 2 | 12 | 23 | 41 |
| 101 | M | -40.6498921368397 | -20.8069069334279 | 2 | 1982 | 4100 | 189 | 12 | 24 | 40 |
| 102 | M | -47.0174640004126 | -24.3839659997340 | 3 | 9624 | 407 | 0 | 12 | 15 | 22 |
| 103 | M | -43.7685686236329 | -22.9328942480597 | 2 | 3354 | 8014 | 9 | 12 | 21 | 44 |
| 104 | M | -43.2315805796622 | -22.7387391311087 | 2 | 4841 | 8197 | 4 | 9 | 9 | 35 |
| 105 | M | -40.1526238168947 | -19.9497258505292 | 2 | 1396 | 1673 | 0 | 9 | 36 | 2 |
| 106 | M | -43.5992298386331 | -23.0255913900612 | 3 | 13252 | 112 | 0 | 9 | 23 | 18 |
| 107 | M | -40.2824531538752 | -20.2761340284942 | 2 | 2933 | 15223 | 2 | 9 | 30 | 44 |
| 108 | M | -43.0327531196711 | -22.6940626351244 | 2 | 41978 | 268 | 0 | 14 | 3 | 10 |
| 109 | M | -46.1782544104794 | -23.9237411528265 | 3 | 2497 | 45227 | 0 | 14 | 21 | 16 |
| 110 | M | -48.6635513311815 | -26.5801677694320 | 3 | 7353 | 794 | 6 | 15 | 19 | 36 |
| 111 | M | -48.6542030288688 | -26.7720904630730 | 3 | 2538 | 7588 | 2250 | 15 | 20 | 42 |
| 112 | M | -48.5128154752146 | -26.2245731344742 | 3 | 4696 | 1691 | 15 | 15 | 20 | 15 |
| 113 | M | -48.5885959471456 | -27.1407648253842 | 3 | 3519 | 498 | 0 | 15 | 21 | 21 |
| 114 | M | -48.7662483390382 | -28.4780005612478 | 3 | 1494 | 2469 | 58 | 15 | 27 | 41 |
| 115 | M | -48.6231323880948 | -27.8176427163186 | 3 | 1570 | 409 | 0 | 15 | 21 | 7 |
| 116 | M | -48.5995578771934 | -26.4606113424470 | 3 | 3612 | 573 | 0 | 15 | 22 | 13 |
| 117 | M | -48.5087521091206 | -27.5475478323507 | 3 | 4656 | 3860 | 117 | 15 | 22 | 41 |
| 118 | M | -48.6509266991050 | -27.6562317173321 | 3 | 1835 | 4413 | 656 | 15 | 22 | 6 |

| | | | | | | | | | | |
|---|---|---|---|---|---|---|---|---|---|---|
| 119 | M | -48.4881454331297 | -27.4373905653624 | 3 | 3080 | 565 | 0 | 15 | 23 | 7 |
| 120 | M | -48.5253426351124 | -27.6497684213402 | 3 | 1630 | 1290 | 33 | 15 | 21 | 15 |
| 121 | M | -48.5676237292353 | -25.8858027255023 | 3 | 13659 | 1408 | 0 | 15 | 21 | 7 |
| 122 | M | -48.5842534612152 | -26.1615518884757 | 3 | 3412 | 12001 | 639 | 15 | 19 | 40 |
| 123 | S | -57.4559774565656 | -37.7853073407575 | 5 | 0 | 2934 | 0 | 15 | NA | 31 |
| 124 | S | -64.4151642291526 | -54.7528812904285 | 5 | 0 | 0 | 0 | NA | NA | NA |
| 125 | S | -62.2665539972735 | -40.2984757657098 | 5 | 0 | 0 | 0 | NA | NA | NA |
| 126 | S | -62.3975376673675 | -38.7956709872570 | 5 | 0 | 0 | 0 | NA | NA | NA |
| 127 | S | -64.9177206595016 | -40.7514905228330 | 5 | 0 | 0 | 0 | NA | NA | NA |
| 128 | S | -64.1090728470970 | -41.0073600867404 | 5 | 0 | 0 | 0 | NA | NA | NA |
| 129 | S | -64.3054492604958 | -42.4049826283180 | 5 | 0 | 0 | 0 | NA | NA | NA |
| 130 | S | -65.9659837603658 | -47.7520672064788 | 5 | 0 | 0 | 0 | NA | NA | NA |
| 131 | S | -63.6769990158912 | -42.2186598971545 | 5 | 0 | 0 | 0 | NA | NA | NA |
| 132 | S | -65.9333607567302 | -47.7505872261499 | 5 | 0 | 0 | 0 | NA | NA | NA |
| 133 | S | -65.9333607567302 | -47.7505872261499 | 5 | 0 | 0 | 0 | NA | NA | NA |
| 134 | S | -68.9732705227977 | -51.4908751001641 | 5 | 0 | 0 | 0 | NA | NA | NA |
| 135 | S | -57.3301021300384 | -37.6690312812071 | 5 | 0 | 0 | 0 | NA | NA | NA |
| 136 | S | -57.3825314343455 | -35.9667610903556 | 5 | 0 | 0 | 0 | NA | NA | NA |
| 137 | S | -63.6375005309851 | -42.7179191582481 | 5 | 0 | 0 | 0 | NA | NA | NA |
| 138 | S | -56.7764408212518 | -36.2975199127133 | 5 | 0 | 0 | 0 | NA | NA | NA |
| 139 | S | -69.1759416574481 | -50.9754967166561 | 5 | 0 | 0 | 0 | NA | NA | NA |
| 140 | S | -68.9732705227977 | -51.4908751001641 | 5 | 0 | 0 | 0 | NA | NA | NA |
| 141 | S | -56.6974548373210 | -36.4365751427255 | 5 | 0 | 0 | 0 | NA | NA | NA |
| 142 | S | -68.6068321566295 | -53.0453699955838 | 5 | 0 | 0 | 0 | NA | NA | NA |
| 143 | S | -48.3985656562632 | -25.6103239615379 | 3 | 0 | 0 | 0 | NA | NA | NA |
| 144 | S | -42.2901583596797 | -22.9389620501455 | 2 | 0 | 3264 | 0 | 15 | NA | 45 |
| 145 | S | -48.3480623452679 | -25.5793893465439 | 3 | 0 | 386 | 0 | 6 | NA | 41 |
| 146 | S | -48.5643609662346 | -25.8994021435012 | 3 | 0 | 457 | 0 | 15 | NA | 34 |
| 147 | S | -50.9177829996883 | -31.2598569988616 | 4 | 0 | 8917 | 0 | 15 | NA | 43 |
| 148 | S | -69.4602956300465 | -52.4691231992791 | 5 | 0 | 38 | 0 | 9 | NA | 14 |
| 149 | S | -69.2061873857517 | -52.6731616152896 | 5 | 0 | 0 | 0 | NA | NA | NA |
| 150 | S | -56.0270659157636 | -34.8800729306977 | 5 | 0 | 0 | 0 | NA | NA | NA |
| 151 | S | -55.7302763449119 | -34.7725380507189 | 5 | 0 | 0 | 0 | NA | NA | NA |
| 152 | S | -55.3842366858992 | -34.8023333687683 | 5 | 0 | 0 | 0 | NA | NA | NA |
| 153 | S | -55.8680502615481 | -34.7971762966657 | 5 | 0 | 0 | 0 | NA | NA | NA |
| 154 | S | -56.3736533791095 | -34.7869899200344 | 5 | 0 | 0 | 0 | NA | NA | NA |
| 155 | S | -54.2506291209729 | -34.6749926267404 | 5 | 0 | 0 | 0 | NA | NA | NA |
| 156 | S | -56.2853924654242 | -34.8966227648919 | 5 | 0 | 0 | 0 | NA | NA | NA |
